# Ultra-sensitive measurement of brain penetration mechanics and blood vessel rupture with microscale probes

**DOI:** 10.1101/2020.09.21.306498

**Authors:** Abdulmalik Obaid, Mina-Elraheb Hanna, Song-Wen Huang, Yu-Ting Hu, Omar Jáidar, William Nix, Jun B. Ding, Nicholas A. Melosh, Yu-Wei Wu

## Abstract

Microscale electrodes, on the order of 10-100 μm, are rapidly becoming critical tools for neuroscience and brain-machine interfaces (BMIs) for their high channel counts and spatial resolution, yet the mechanical details of how probes at this scale insert into brain tissue are largely unknown. Here, we performed quantitative measurements of the force and compression mechanics together with real-time microscopy for *in vivo* insertion of a systematic series of microelectrode probes as a function of diameter (7.5–100 μm and rectangular Neuropixels) and tip geometry (flat, angled, and electrochemically sharpened). Results elucidated the role of tip geometry, surface forces, and mechanical scaling with diameter. Surprisingly, the insertion force post-pia penetration was constant with distance and did not depend on tip shape. Real-time microscopy revealed that at small enough lengthscales (<25 μm), blood vessel rupture and bleeding during implantation could be entirely avoided. This appears to occur via vessel displacement, avoiding capture on the probe surface which led to elongation and tearing for larger probes. We propose a new, three-zone model to account for the probe size dependence of bleeding, and provide mechanistic guidance for probe design.

**Significance Statement:** Microscale neural probes are central to next-generation brain–machine interfaces, yet how they physically penetrate living brain remains poorly quantified. Using a high-sensitivity force sensor integrated with real-time microscopy, we measured *in vivo* force–displacement and visualized vascular responses for microwires (7.5–100 μm) and Neuropixels. We find that once the brain’s protective pia membrane is breached, insertion force remains essentially constant with depth, while pia puncture force and pre-penetration compression scale linearly with probe diameter. Real-time imaging reveals a sub-25 μm regime in which blood vessels are displaced rather than ruptured. These results motivate a three-zone model of vessel capture versus displacement and provide actionable mechanical design rules for low-trauma, high-density neural interfaces.

## Introduction

Microelectrodes implanted into the brain are a critical component of new neuroprosthetic applications and brain-machine interfaces (BMIs), including high-density silicon probes (70 x 20 μm)(1, 2), syringe injectable electronics (∼100 μm) (3), shuttle delivery (4), and microwire arrays (<20 μm per wire) (5-9). As these new probes become more widespread and new fabrication techniques become available, understanding the mechanics of brain penetration and insertion of devices in the 10-100 μm size scale is essential for balancing geometric design, materials strength and tissue damage (10-12). Appropriate optimization of mechanical insertion into the tissue is critical for success, as large devices can cause traumatic tissue damage and scar formation, while thin devices may buckle under the loads necessary to penetrate the outer pia membrane (5, 13-16).

Here, we performed systematic measurements of *in vivo* brain insertion forces and tissue compression for microscale probes of different diameters (7.5–100 μm cylindrical microwires and rectangular Neuropixels) and tip geometry to provide a rigorous mechanical framework for guiding probe design and understanding in this critical size regime. Microwires are a useful model system in that they are available in a wide variety of sizes with uniform surfaces and tip shapes, and have long been used for recording structures deep in the brain (13, 15, 17-19). While the general expectation is that smaller wire diameters should yield lower damage, actual data for insertion mechanics is sparse. Prior investigations in the 10–100 μm probe size range have been limited to a few different sizes with relatively low sensitivity measurement units. Initial tissue damage studies with lower-resolution instrumentation were inconclusive about size dependence, with no significant differences found between large (cross-sectional dimensions ∼ 200 x 60 μm) and small (100-140 x 15 μm) devices (20).

Measuring the mechanics of such ultra-small devices inserting into the brain, a soft, ultra-compliant material, are highly challenging due to the large insertion displacements (mm to cm) before penetration, together with rapid and/or small force events present during insertion. Compromises typically have to be made either in force sensitivity or temporal resolution; many features may have been missed in previous studies simply due to instrumental limitations (13, 21-23).

To address these challenges, we developed a high-performance force-displacement measurement system using a modified nanoindentation head as a force transducer. Nanoindentation transducers are among the fastest, most sensitive force-displacement systems available with 3 nN force and 1 ms temporal sensitivity. However, these instruments are designed with maximum displacements on the order of hundreds of microns (24), compared to the millimeters of compression required before tissue penetration. This large displacement range was achieved by integrating an entire *iNano* nanoindenter (Nanomechanics, Inc.) measurement head onto a long-travel linear actuator, and measuring force and time with the nanoindenter in a zero-displacement mode while the actuator traveled at a constant velocity.

In combination with the mechanical measurements we visualized the probe inserting with real-time epifluorescence or two-photon microscopy. These measurements allowed direct visualization of how blood vessels failed during insertion, leading to the discovery of a probe size regime where vessel rupture can be avoided altogether. We propose a new three-zone model for blood vessel rupture to explain these observations, which predicts that sufficiently small probes will displace blood vessels.

## Results

### Penetration force measurement

A high-performance mechanical measurement system was developed to monitor the forces and displacements during penetration of soft (∼1 kPa) brain tissue and mimics. This apparatus (**Fig. 1a**) used a *NanoMechanics iNano InForce 50* indentation head as the force transducer. The transducer position relative to the tissue is then controlled via a low-noise linear actuator moving 20 μm/s to a depth of 2.5 mm. Tungsten microwire diameters of 7.5, 15, 25, 35, 50, 80, and 100 μm were measured, each with smooth sides. Different tip shapes (flat polished, angle polished, and electrosharpened) and tissue systems (phantom, *ex vivo* and *in vivo*) were prepared as described in the Methods section.

**Figure 1.**
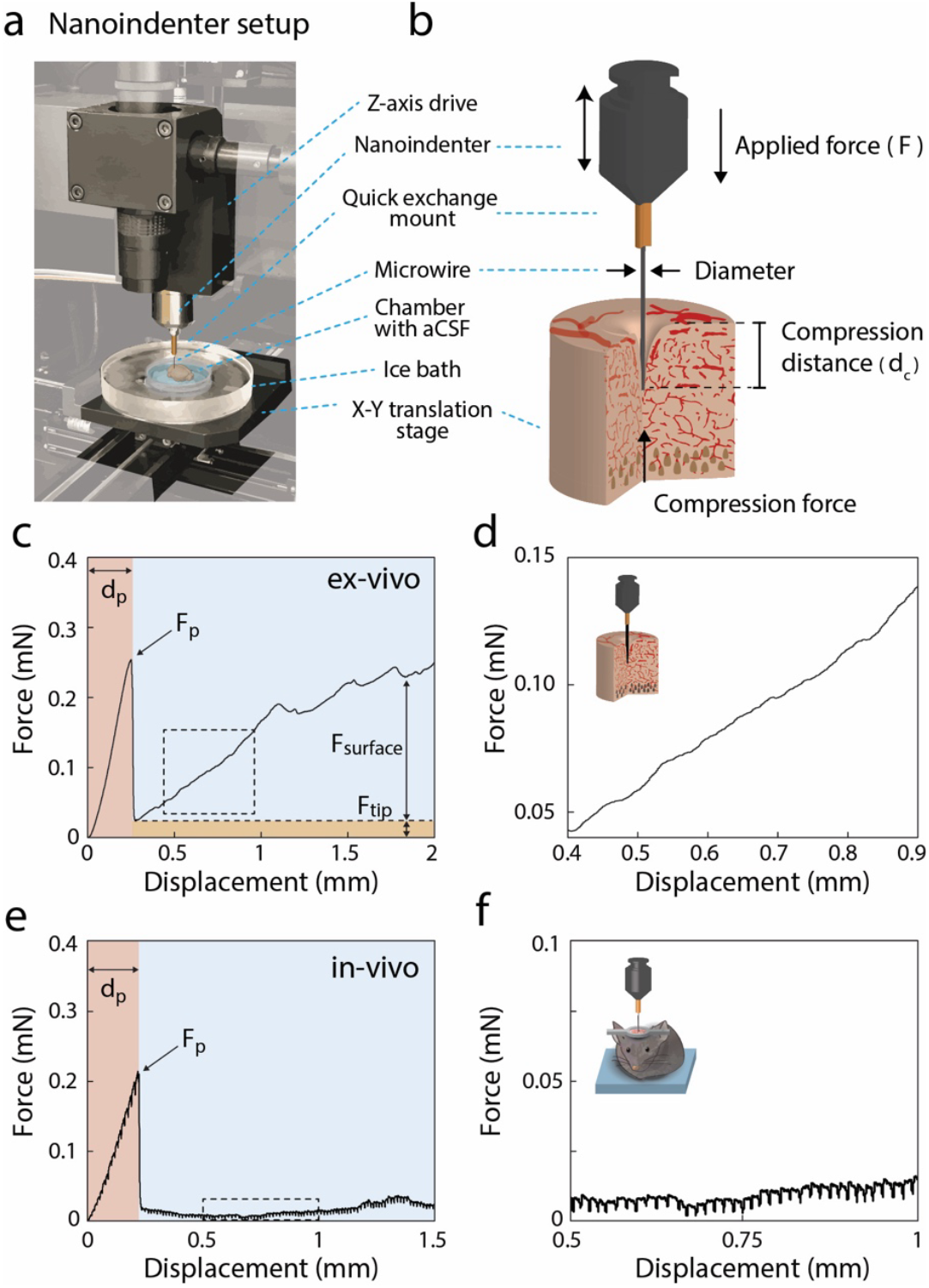
Penetration of brain mimics, *ex vivo* and *in vivo* brain tissue. **(a)** Image of the adapted nanoindenter setup. Background electronics are grayed out to highlight areas of interest. The nanoindenter is mounted on a z-axis drive that provides the vertical translation. **(b)** Schematic illustration of insertion into brain tissue. **(c,d)** Force-displacement curve for insertion of 15 μm diameter flat-polished wire into acutely excised brain tissue. The force increases exponentially pre-penetration, then increases linearly as a function of depth post-penetration. The linear increase in force is attributed to surfaces forces along the length of the wire (*F*_*surface*_) in addition to the constant force at the tip (*F*_*tip*_). **(e,f)** Force-displacement curve for insertion of 15 μm diameter flat-polished wires *in vivo*. Insertion into live brain tissue also displays a sharp drop in force during penetration, but the force plateaus as the wire is driven deeper, unlike the *ex vivo* case. Oscillations during insertion into live brain are consistent with breathing and heart rate.

A schematic of the experimental apparatus and representative force-extension curves for both freshly excised *ex-vivo* (**Fig. 1c,d**) and *in vivo* samples (**Fig. 1e,f**). As a 15 μm microwire extended at 20 μm/s, the force increased exponentially for the first ∼250 μm, roughly ∼15 times the diameter of the wire. Visually, as the wire is displaced the brain dimples without penetration, until a critical force is reached and the probe penetrates through the pia and a sudden drop in force is observed. This corresponded with the tissue visually relaxing around the wire, with the surface approaching its original location, which we interpret as indicating the microwire was inside the brain. From this penetration event both the force at puncture (*F*_*p*_) and the displacement to puncture (*d*_*p*_) were recorded.

In a subset of control experiments, an alternative measurement configuration using a *MTS Nano Indenter XP* was employed. In this setup, the indenter transducer position was fixed while a low-noise linear actuator advanced the sample upward at velocities of 5 and 20 μm s−^1^ to depths of ∼2 mm (**Fig. S1a,b**). To verify that the measured puncture forces reflected pia penetration rather than incomplete removal of the dura mater, we performed additional insertions through intact, moist dura prior to dura removal (**Fig. S1c–f**). Dura removal was confirmed visually under a dissection microscope and independently verified using two-photon microscopy with second-harmonic generation (SHG) imaging (**Fig. S1c,d**). Insertions through intact dura required substantially larger *Fp* (>10 times lager) and exhibited prolonged loading phases (*d*_*p*_) consistent with the higher stiffness of the dura layer (**Fig. S1e,f**). Following surgical removal of the dura, puncture forces decreased markedly, and force–displacement traces adopted the characteristic signatures observed throughout this study. Together, these controls confirm that the forces reported here reflect pia penetration mechanics, rather than the composite mechanics of the dura–arachnoid–pia complex.

Note that the force oscillations *in vivo* are not noise or stick-slip events, but instead the pulsatile effects from heartbeat and breathing (**Fig. 1f**). Both the curve shape and magnitude of the pre-penetration phase are very similar between *in vivo* and *ex vivo* tests, which agrees with previous indentation measurements suggesting they have similar mechanical properties (25-27).

Surprisingly, as the insertion force to drive the wire deeper into tissue *in vivo* after pia-penetration was constant as a function of depth. Representative force–displacement traces across wire diameters illustrating this depth-independent post-penetration force are shown in **Fig. S6**. Slight features or humps are observed in the force–displacement curves, but without the monotonic linear force increases observed *ex vivo*. It is surprising that there is no increased resistance to insertion despite more probe area was in contact with the brain, yet this trend was highly consistent over dozens of experiments. We hypothesize this may be due to lubrication from active pumping of cerebrospinal fluid (CSF) or the dynamic motion from heartbeats preventing surface adhesion or static friction, yet more study needs to be done.

The absence of a linear increase in force post pia puncture *in vivo* shows that the pia is the primary mechanical barrier to electrode insertion; if a probe is stiff enough to penetrate the pia, it can be inserted to arbitrary depths. This suggests that initial penetration into brain tissue is the critical mechanical challenge in designing penetrating electrodes. Once implants are past the pia, there is minimal mechanical barrier to drive deeper into tissue; the force to insert 500 μm deep is very similar to the force to insert two millimeters.

These penetration results differed markedly from those obtained in agarose phantoms, despite matching the bulk elastic modulus of brain tissue. For a 0.6% agarose gel (**Fig. S2a-d**), penetration produced pronounced saw-tooth features in the force–displacement curves at both micrometer and millimeter scales, behavior not observed in either *ex vivo* or *in vivo* brain. Although agarose hydrogels at this concentration are commonly used as mechanical brain mimics for indentation studies (22, 23, 28), their penetration response reflects fracture- and adhesion-dominated mechanics rather than the smooth insertion dynamics seen in living tissue. Reducing the concentration to 0.3% agarose resulted in fewer saw-tooth features following initial penetration and substantially lower *F*_*p*_ **(Fig. S2e–f)**, yet the overall insertion behavior remained qualitatively distinct from brain tissue. Thus, while agarose gels are useful for approximating bulk elastic properties, they do not faithfully reproduce the mechanics of probe penetration and are not suitable surrogates for penetration studies.

### Influence of Microwire Diameter

To understand the role of electrode size on brain insertion mechanics, the force-displacement to insert wires with diameters between 7.5 μm to 100 μm was measured *ex vivo* and *in vivo* (**Fig. 2**). All of these microwires had highly consistent, flat-tipped geometries by polishing the distal ends; tip shape dependence will be discussed below. The slope of the initial loading, indicating the compliance of the brain tissue, was only weakly dependent on wire diameter, with slightly lower compliances for smaller probes (**Fig. S3**). This suggests that over the range of the wire diameters studied, bulk brain tissue mechanics are generally homogeneous, consistent with previous studies on indentation into brain tissue (24, 29), and our measurements were insensitive to the order of wire diameters tested across animals (**Fig. S5**).

**Figure 2.**
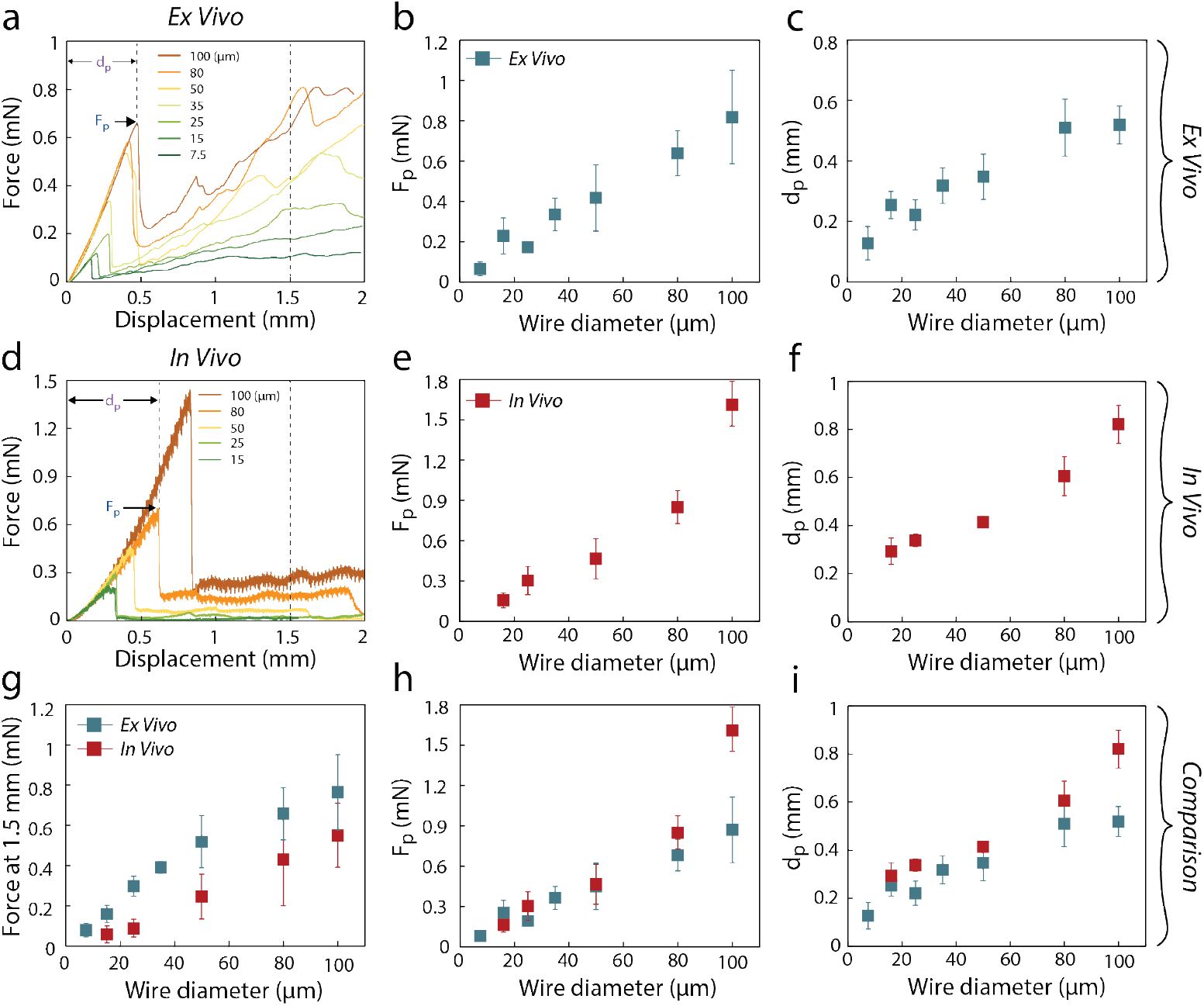
Influence of microwire diameter. **(a)** Representative force-displacement curves comparing the response of different diameter flat-polished probes inserted into *ex vivo* brain at a rate of 20 μm/s. Pia puncture occurs during abrupt drop in measured force, wherein the puncture force (*F*_*p*_) and displacement to puncture (*d*_*p*_) are extracted. **(b)** Relationship between puncture force and microwire is found to be linear. **(c)** The displacement to puncture as a function of wire diameter is also linear. **(d)** Representative force-displacement curves of flat-polished probes of increasing diameter inserted into live brain tissue at 20 μm/s. **(e,f)** Relationship between puncture force and displacement to puncture as a function of size for insertion into live brain. **(g)** The force at 1.5 mm as a function of wire diameter for flat-polished wires, comparing *ex vivo* and *in vivo*. The forces after penetration are significantly lower *in vivo* than from *ex vivo* insertions. **(h,i)** Comparison of the scaling of puncture force and displacement to puncture with size between *ex vivo* and *in vivo*. Forces and displacements are found to be similar 50 μm and below, with larger differences observed *in vivo* at large sizes. Data are shown as mean ± SD. For each diameter, measurements were obtained from *ex vivo*: n = 3 (7.5 μm), 7 (15 μm), 5 (25 μm), 4 (35 μm), 4 (50 μm), 5 (80 μm), and 5 (100 μm) independent insertions of each wire size, pooled across N = 4 mice; *in vivo*: n = 4 (15 μm), 5 (25 μm), 5 (50 μm), 4 (80 μm), and 4 (100 μm) independent insertions of each wire size, pooled across N = 5 mice (multiple insertions per animal, spaced ≥500 μm apart). Each insertion was treated as an independent mechanical trial.

The pia penetration event shape was highly stereotyped across all sizes, yet the force magnitudes to cause puncture (*F*_*p*_) varied by over an order of magnitude; from 185 ± 40 μN for 15 μm wires, to 1610 ± 239 μN for 100 μm diameter *in vivo*. Both the *F*_*p*_ and compression *d*_*p*_ scaled linearly with probe diameter, rather than the cross-sectional area of the tip. This observation is consistent with models of crack-initiated or energy-limited failure in compliant materials (30, 31), indicating the failure of the pia may occur via one of these modes. From this data, the force and compression needed to puncture the brain could be directly calculated based on the size of the wire *in vivo*:

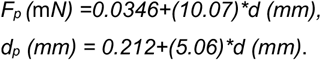

The insertion force past pia penetration was roughly constant for all *in vivo* tests (**Fig. 2d**), yet the magnitude of this force scaled linearly with diameter (**Fig. 2e**). Consistent with recent *in vivo* demonstrations of bleeding-absent insertions using anisotropically relaxing polymer probes, reducing the effective cross-section shifts the failure mode from vascular penetration to vessel deflection, lowering acute hemorrhage risk (32). Larger probes required larger absolute magnitudes of insertion force, yet did not increase significantly once through the pia. These observations are consistent with a friction model for *in vivo* tissue insertion, where the normal restoring force arises from lateral tissue displacement proportional to the probe’s diameter. The linear relationship suggests the coefficient of friction is constant over all probe diameters studied. Interestingly, a significant difference between *ex vivo* and *in vivo* penetration force and compression exists for microwires of 100 μm diameter (p < 0.05, two-way ANOVA, *post hoc* Bonferroni) (**Fig. 2h,i**). The significantly higher forces required for penetration *in vivo* for larger wires is unclear at the moment, yet may involve pressurization of the vasculature due to blood flow (33), or continual tissue movement due to breathing and heartbeat.

### Tip geometry dependence

Sharpening the probe tip geometry is a common strategy for reducing tissue compression in penetrating microelectrodes (34, 35) which may result in less tissue damage, such as for Michigan-style probe arrays (36, 37). To elucidate the effects of probe shape in the 10-100 μm range, penetration of angle-polished (24º) and electrosharpened probes (tip radius ∼10 nm, **Fig. 3a,b**) were studied in comparison to the flat-polished tips. Two representative force-displacement curves for 15 μm and 80 μm wires of the three different tip geometries are shown in **Fig. 3c** and **3d** for *in vivo* tests (*ex vivo* shown in **Fig. S4**). Flat- and angle-polished wires showed no statistically different behavior for microwires ≤ 100 μm in diameter (**Fig. 3e**; displacement scaling shown in **Fig. S7**), indicating at this lengthscale angle sharpening has no effect. Yet the results diverged >100 μm, where the force and distance for angle-polished tips were lower than flat-polished tips. To further confirm this trend, microwires with 125 μm diameter were tested, and *Fp* were again lower for angle-polished tips, in agreement with results for millimeter-scale needles (21, 38). This data suggests that our intuitive expectation that angled ‘sharp’ tips are better is true at macroscopic lengthscales, but breaks down in the <100 μm regime.

**Figure 3.**
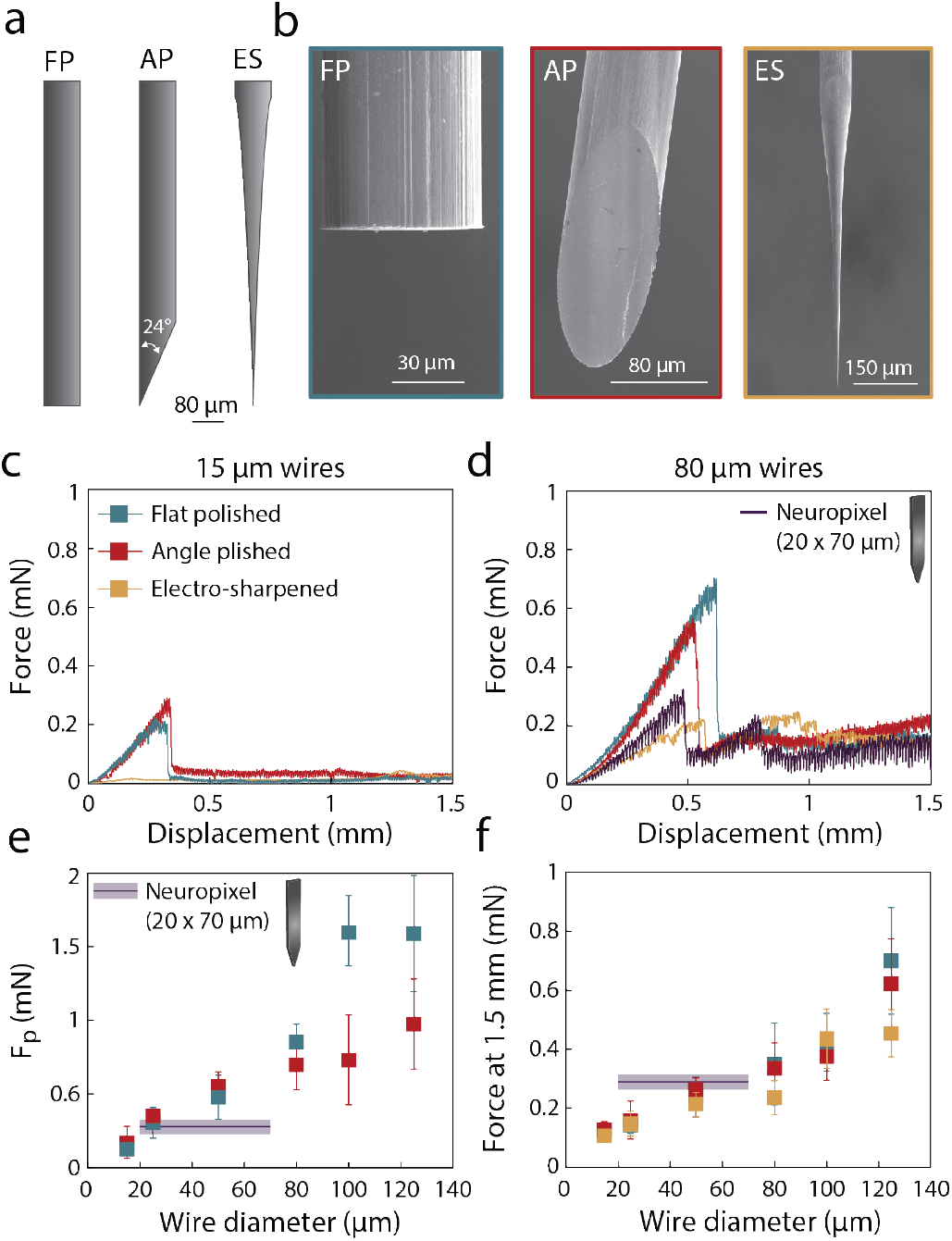
Tip geometry dependence. **(a)** Graphical illustration of the three different tip geometries studied; flat-polished (FP), angle-polished (AP), and electrosharpened (ES). **(b)** Representative SEM images of the three different tip geometries. **(c,d)** *In vivo* force-displacement curves for 3 tip geometries for 15 μm diameter **(c)** and 80 μm diameter **(d)** microwires. A silicon probe (*Neuropixels*) with a 70 μm wide, 20 μm thick geometry is plotted in comparison with 80 μm wires **(d). (e)** Measured puncture force *in vivo* as a function of wire diameter for FP and AP wires and a *Neuropixels. Neuropixels* values are plotted with wire diameter values between 20 and 70 μm because of their rectangular cross-section. **(f)** *In vivo* force at 1.5 mm depth as a function of wire diameter for all tip geometries tested, including *Neuropixels*. Data are shown as mean ± SD. For each diameter, measurements were obtained from FP: n = 4 (15 μm), 5 (25 μm), 5 (50 μm), 4 (80 μm), 4 (100 μm), 3 (125 μm); AP: n = 4 (all sizes) and EP: 4 (15 μm), 4 (25 μm), 3 (50 μm), 4 (80 μm), 3 (100 μm), 4 (125 μm) independent insertions of each wire size, pooled across N = 5 mice; (multiple insertions per animal, spaced ≥500 μm apart). Each insertion was treated as an independent mechanical trial.

In contrast, electrosharpened wires displayed strikingly different behavior, with no discernable pia penetration event (**Fig. 3c,d**). There are numerous small rapid rises and drops in the force readout during insertion, but none could be classified as a distinct penetration of the brain surface. Synchronized videos of electrosharpened wire insertion also show no detectable insertion event, dimpling or tissue relaxation (**Video S1**). Hence, we cannot report a puncture force for electrosharpened wires. The forces during initial insertion into brain tissue were roughly an order of magnitude smaller than flat tips, ranging between 10 to 100 μN. Empirically, an electrosharpened tip that has been dulled or bent will still result in a distinguishable puncture event, albeit of much lower peak force magnitude than flat or angle polished microwires.

Interestingly, after pia puncture no insertion force difference was observed between flat, angled, or electrosharpened tips (**Fig. 3f**). This reveals that after penetration, the insertion force is dominated by surface forces along the shaft of the electrode and the surrounding brain tissue, rather than effects at the probe tip.

### Comparison to rectangular *Neuropixels* probes

With the increasing prevalence of Michigan-style and *Neuropixels* silicon probes for high-density neural recording(1, 2), we measured the mechanics of inserting rectangular 20 x 70 μm cross-section *Neuropixels* probes (**Fig. 3e,f** and **Fig. S8**). *Neuropixels* required pia penetration forces similar to that of 25 μm diameter wires, which is close to the 20 μm thickness of the probe tip (**Fig. 3e**). Past pia penetration, the insertion forces corresponded to cylindrical wires with diameters between 50 μm and 80 μm (**Fig. 3f**). *Neuropixels* have an equivalent surface area of a ∼57.5 μm diameter wire, thus the insertion force appears dependent on circumference or surface area, rather than cross-sectional shape. These results are commensurate with the force independence on tip shape, indicating surface forces along the shaft of the probe dominate the bulk tissue insertion force, regardless of geometry. The effect of insertion speed was also studied using the *Neuropixels*, inserting at 2, 20 and 100 μm/s (**Fig. S8g**). While the displacement at puncture remained similar across speeds, the force magnitude (both peak and post-penetration plateau) increased with insertion speed, consistent with the viscoelastic properties of brain tissue (39). Finally, *Neuropixels* often exhibited a secondary ‘peak’ in the force curve after the pia penetration (**Fig. 3d**, purple trace). The origin of this event is unclear, but was quite common, observed in ∼80% of the force traces. To assess whether insertion speed modulates these diameter-dependent relationships, we performed additional *in vivo* insertions at a slower speed (5 μm/s) using 20 μm and 80 μm flat-polished microwires (**Fig. S8 h,i**). Compared to insertions at 20 μm/s, slower insertions slightly increased *F*_*p*_ and *dp* at 5 μm/s for either diameter, and post-penetration forces did not show clear changes with depth. These results indicate that within the range tested, only slightly alters the scaling of penetration mechanics with probe diameter.

### Effect of probe size on bleeding

Bleeding during or post probe-insertion is a strong indicator of vascular and/or tissue damage, and can lead to serious subsequent trauma such as vasospasms. Larger probes, such as deep brain stimulation electrodes, always involve some level of hemorrhaging, yet probes in the sub-100 μm range have already shown different mechanical insertion behavior, thus may have different vessel damage as well. To quantify a bleeding event, we carefully observed with a magnified camera any surface bleeding, either during insertion or after the probe was withdrawn from the tissue (**Fig 4a,b**). These events were tallied on a binary yes/no basis, as the magnitude of hemorrhaging was difficult to accurately quantify.

**Figure 4.**
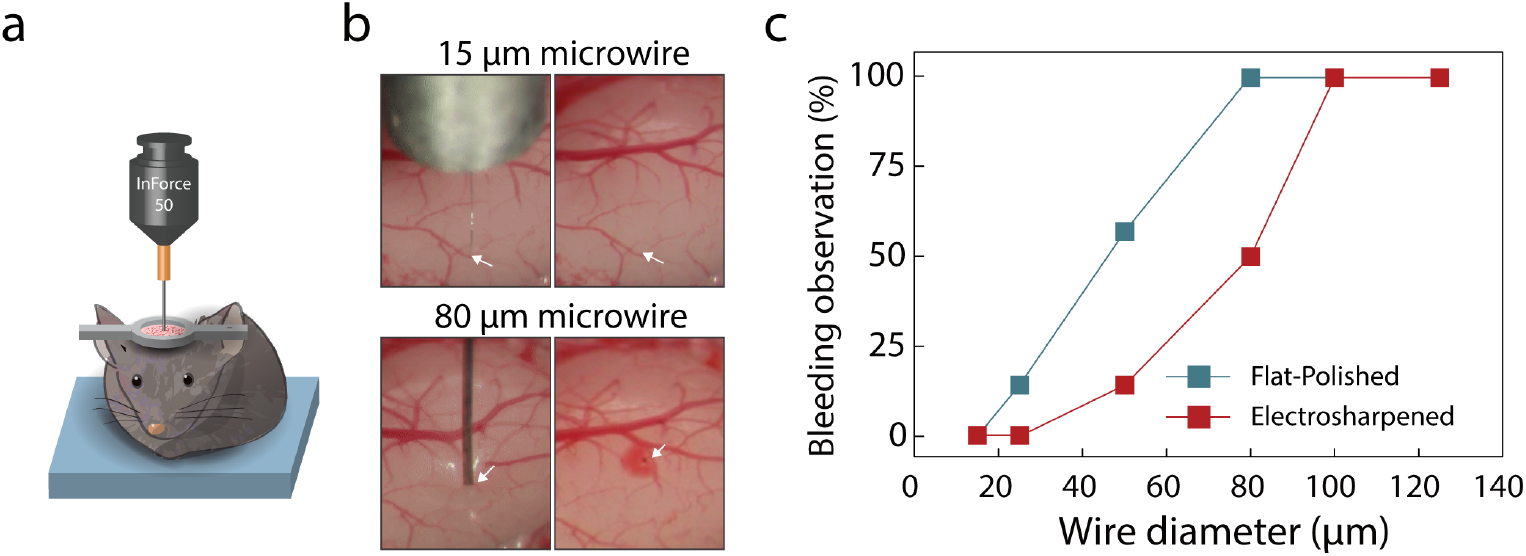
Surface bleeding observations. **(a)** Schematic of the *in vivo* force measurements. **(b)** Representative images of the surface of the brain before (*left*) and after (*right*) insertion of 15 μm and 80 μm microwires. No bleeding is observed is on the surface of the brain for 15 μm wires. After retraction of the 80 μm wire, the insertion site is marked by blood leaking from the surface. **(c)** Percent bleeding observed optically at the surface of the brain for insertion of flat-polished and electrosharpened wires. For insertion of 15 μm wires, no bleeding was observed for all tip geometries. Between 25 μm and 80 μm, electrosharpened wires showed fewer incidence of bleeding at the surface in comparison to flat-polished tips. For > 100 μm, all wires displayed bleeding at the surface. Large, visible vessels were avoided for all measurements. Bleeding observation was obtained from FP: n = 5 (15 μm), 7 (25 μm), 12 (50 μm), 5 (80 μm), 5 (100 μm), 3 (125 μm) and EP: 5 (15 μm), 7 (25 μm), 7 (50 μm), 4 (80 μm), 5 (100 μm), 3 (125 μm) independent insertions of each wire size, pooled across N = 5 mice; (multiple insertions per animal, spaced ≥500 μm apart).

The results clearly show that the incidence of bleeding is strongly correlated with probe size, with a roughly sigmoidal dependence (**Fig. 4c**). From these observations, for ≥ 100 μm probes there is always bleeding, transitioning to less common bleeding events between 100 to 25 μm diameters, and finally to where no bleeding occurs for probes < 25 μm. Interestingly, electrosharpening the tip produces an offset in the curve rather than changing its overall shape, reducing the ‘effective’ wire diameter by 20–30 μm (**Fig. 4c**, red trace). Thus, bleeding can be reduced by selecting sharper tips, yet probe diameter is still the most important variable for hemorrhaging. For example, insertions for all 100 μm tip shapes caused bleeding, but no bleeding for 15 μm wires of any tip geometry. These results are highly surprising, suggesting it is possible to completely avoid vascular damage by going to small enough probe lengthscales.

### Monitoring insertion and blood vessel rupture with two-photon imaging and simultaneous force/epifluorescence measurements

To understand probe-tissue interaction during insertion and why bleeding may or may not occur, we modified the apparatus to also perform two-photon microscopy and real-time epifluorescence imaging during microwire insertion simultaneously with force (**Fig. 5a**). These experiments were all performed *in vivo* with either 13, 25 or 80 μm wires with a tail injection of a fluorescent dye (Rhodamine B, **Fig. 5b**) to aid the visualization of the vasculature network and bleeding events.

**Figure 5.**
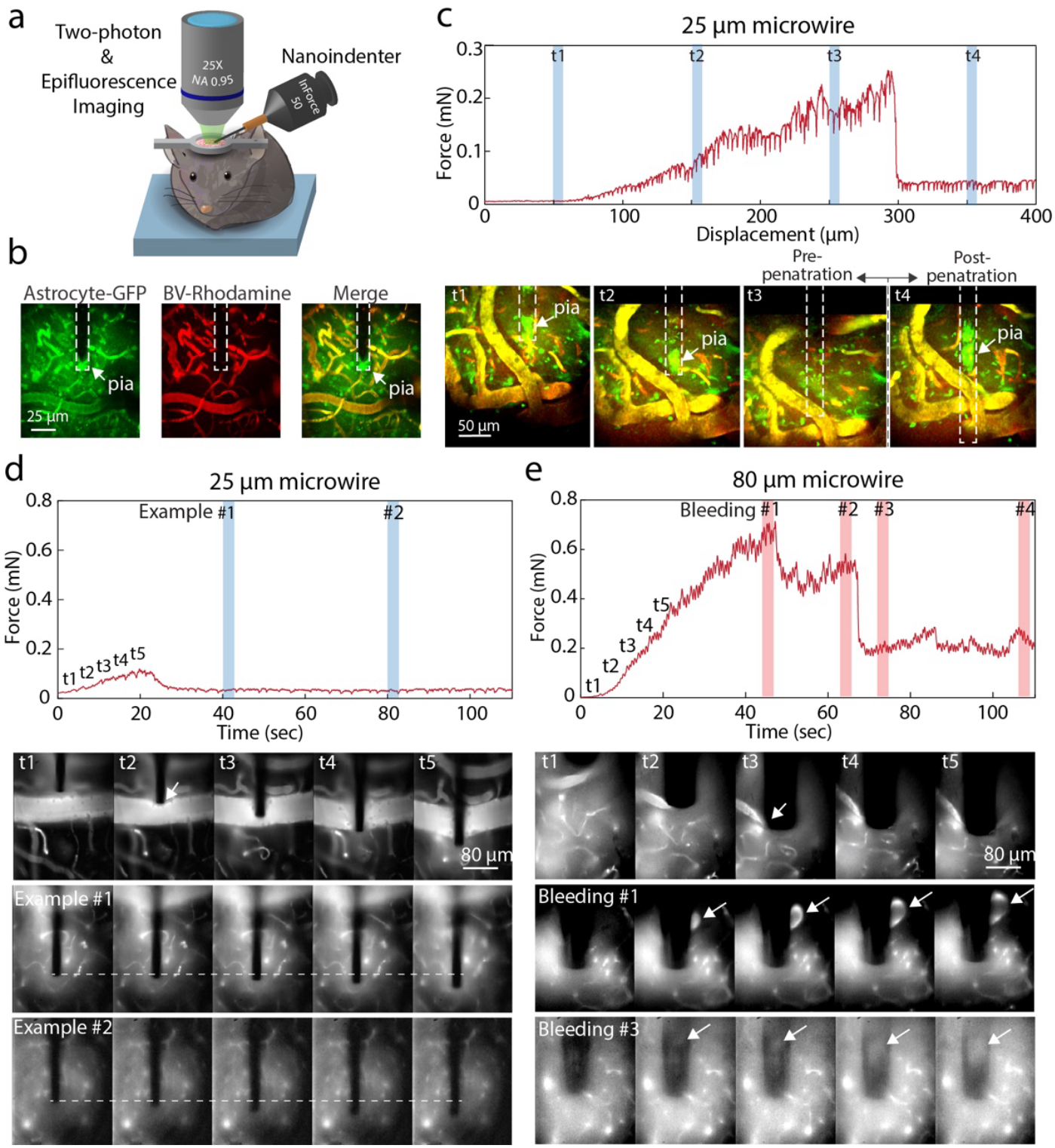
Live two-photon and epifluorescence imaging with *in situ* force measurements. **(a)** Schematic of *in vivo* imaging with *in situ* force measurements. **(b)** eGFP-expressing astrocytes (Astrocyte-GFP) and blood vessels filled via tail vein injection of Rhodamine B (BV-Rhodamine) were imaged with two-photon microscopy. Image stacks were taken with sequential motion of the microwire. **(c)** Force vs displacement during the insertion of a 25 μm microwire and the respective two-photon images at different steps in the insertion process. We observe the exponential increase in force during compression of the surface (*t1-t3)* as well as the lack of increase in force past penetration *(t4)*. Penetration was conformed with 2P imaging, showing the probe is past the pia at *t3,4*. **(d,e)** Force-distance is plotted with synchronized epifluorescence videos of 25 μm and 80 μm microwire insertion, respectively. The exponential increase in force pre-penetration is correlated with the dimpling of the pia and surface vessels around the wire. During insertion of a small wire (25 μm, **c** and **Video S3**), there is no noticeable bleeding or disruption of the vessels, even when the wire comes in contact with microvasculature. In contrast, the 80 μm wire causes multiple bleeding events, both during the initial rupture of the pia surface and vessels beneath the surface.

Forces and tissue displacement were correlated by measuring force vs depth for a 25 μm wire, pausing at certain locations to acquire two-photon (2P) image stacks (**Fig. 5b,c**). No discernible changes were observed in the force measurements during the pauses to acquire the 2P image, suggesting the measurement technique did not alter normal insertion behavior. The location of the pia was identified by a sheath of astrocytes at the surface that is dragged deeper into the brain by the microwire insertion (**Fig. 5b**)(40). Figure 5c shows increasing force pre-pia penetration correlated with dimpling of the pia (visible at the tip of the probe) and compression of surface vessels/tissue around the wire (**Fig. 5c**: *t1* vs *t2*). While the entire tissue appeared to displace vertically, only slight changes in the relative vasculature arrangement were observed. At a critical depth pia rupture occurred, which correlated most notably with the surrounding tissue relaxing vertically (**Fig. 5c**: *t3* vs *t4*). The pia was located mid-way along the microwire, rather than at the tip. In this image stack, no bleeding was observed at any depth.

We then took simultaneous real-time measurements of force and epifluorescence. During insertion of a 25 μm wire at 20 μm/s (**Fig. 5d, Video S2**), there was no noticeable bleeding or disruption of the vasculature, despite the fact that in some cases the probe directly impacted a blood vessel. Timepoints *t1-t5* in **Fig. 5d** show the probe push aside a ∼80 μm blood vessel as the pia is compressed, yet no bleeding occurred even after pia penetration or deeper insertion. The subsequent timespan shows further insertion which also did not disrupt microvasculature in bulk tissue. Instead, this 25 μm probe appeared to deflect the blood vessels to the side as it contacted them, without causing rupture.

This is in stark contrast to a large wire (80 μm, **Fig. 5e** and **Video S3**), wherein significant disruption and bleeding is observed both at and post-pia penetration. During initial force loading pre-penetration (timepoints *t1-t5*), the tissue is compressed and blood vessels were collapsed, but no visible bleeding occurred. Upon partial pia rupture the first bleeding event is visible at the surface, *#1*, and a hemorrhage is observed deeper in the tissue. Additional bleeding events *#2-4* during further insertion were also observed, highlighted in red.

The videos show that blood vessel failure occurred by blood vessels getting trapped on the probe tip surface, then stretching as the probe was inserted further, then finally failing after significant elongation. Note that bleeding event *#1* in Figure 5e occurred by tearing the vessel at a location along the side of probe shaft, rather than at the tip. Thus, vessel rupture did not occur upon initial contact with the probe, nor likely due to a crack propagating near the tip, but only after being caught on the surface of the probe and stretched to failure. These rupture events often corresponded with features in the force profile, such as the sawtooth force profile for bleeding event #4 in Figure 5e. Next, we assessed the effect of insertion on the astrocytic populations and subpial vasculature with two-photon microscopy (**Fig. S9, Video S4**). A 13 μm diameter microwire shows no disruption of the astrocytic populations or vasculature (blood brain barrier) during insertion, akin to the 25 μm diameter microwire, while the 80 μm microwire caused significant disruption of the local vasculature and astrocytic population. After retraction, a clear track of displaced astrocytes and bleeding vessels is left in place of the wire, indicating these larger probes caused significant disruption.

## Discussion

### Three-zone blood vessel rupture model

The results show that smaller electrodes reduce rupture force, tissue compression, and likelihood of vessel rupture. Perhaps most interestingly, we observed a probe size range <25 μm that elicited no bleeding response. From video microscopy, these size wires pushed aside capillaries upon tip approach, allowing passage without causing rupture. For larger probes, blood vessel failure occurred by the vessels becoming captured at or near the probe tip, elongating with further insertion, and finally tearing or rupturing.

From these observations, we propose a three-zone conceptual model for the mechanism for blood vessel rupture (**Fig. 6a**). The model consists of a cylindrical penetrating probe of diameter *D* encountering a blood vessel at some depth. Depending on the location of the vessel and the size of the probe, three distinct events may occur: (1) For vessels in the *capture zone* (red) located underneath the probe tip, blood vessels become entrapped on the surface of the probe, and are stretched as the probe continues to insert. The vessel finally fails due to the strain of elongation, rather than a cutting process near the tip, which often results in a vessel tear along the side of the probe, not underneath the probe itself. (2) In the *displacement zone*, vessels located in a thin ring near the edge of the probe will instead be pushed aside, out from under the probe itself, avoiding capture on the surface. We speculate this occurs due to the differential lateral pressure near the edge of the probe tip, and the width of this zone is roughly constant, as it is a feature of the edge itself rather than the overall probe size. (3) Vessels in the *deformation zone* are compressed and distorted, but do not move significantly relative to the surrounding tissue.

**Figure 6.**
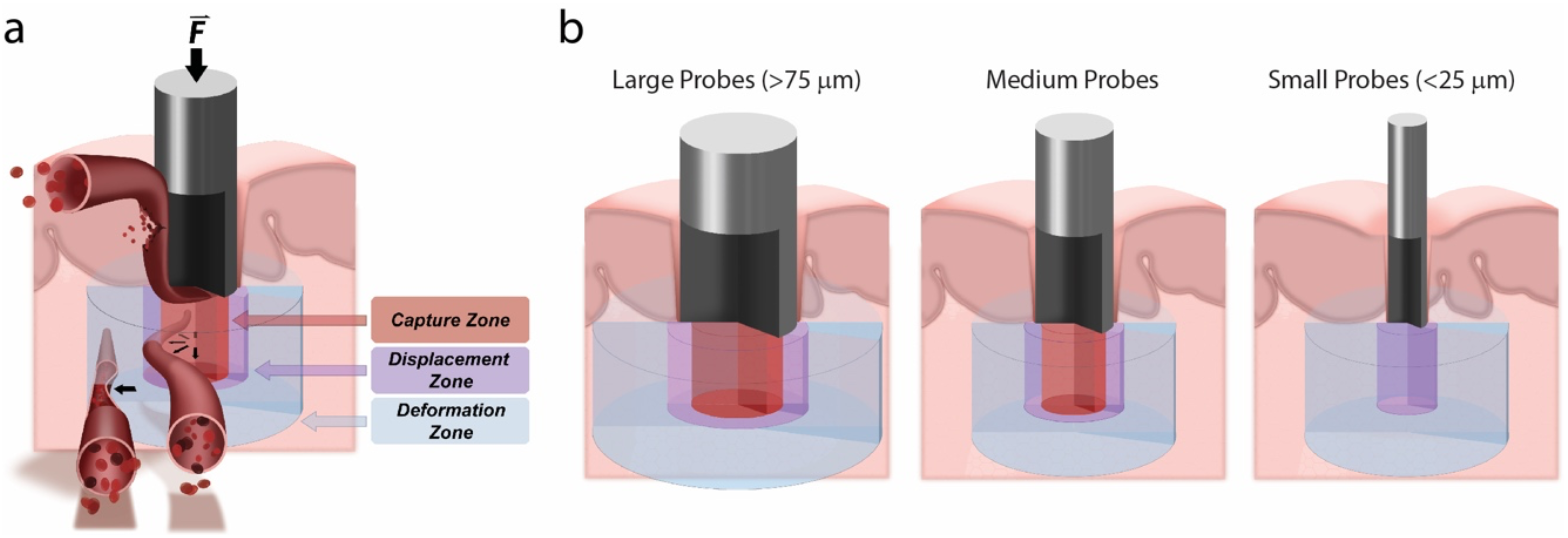
Three zone blood vessel rupture model. **(a)** The three-zone model for blood rupture mechanics consists of a penetrating probe encountering a blood vessel at some depth within the tissue. Depending on the distance of the vessel and the size of the probe, three distinct events may occur: capture and eventual failure, displacement from under the tip, or deformation away from the probe. **(b)** In the proposed model, the size of the displacement zone is roughly constant with diameter, such that at a critical size regime approximately two times the width of the displacement zone, no blood vessel rupture will occur.

We believe this model captures many of the observations made from the force and video evidence and particularly the scaling with probe size (**Fig. 6b**). Large probes would have relatively little lateral displacement force underneath them, thus most of the region beneath the tip is a capture zone, with a thin band of displacement near the edge. As the probe size shrinks the displacement zone stays roughly constant, while the capture zone contracts. Finally, for a sufficiently small probe all vessels are displaced, rather than captured, allowing the probe to be inserted deeply into tissue without damaging the vasculature. Note that a small probe inserted directly on top of a very large blood vessel would still likely eventually cause penetration and bleeding, thus avoidance of large vessels should still be preferred.

These observations also open an interesting insight into how insertion velocity is important at different size scales. Multiple studies have found that injecting large probes very quickly (>1 m/s) could reduce the amount of bleeding and tissue damage, ascribed to the stiffer nature of the viscoelastic tissue at these speeds. However, for the very small <20 μm probes which rely upon vessel displacement, it may be more beneficial to insert slowly (41), allowing time for the vessels to move out of the way. Further velocity measurements are underway, which may shed more light on the insertion mechanics for these small probe dimensions.

## Conclusion

The mechanics of inserting a foreign object into the brain is one of the critical issues for *in vivo* implantations. While numerous studies exist for clinical insertion of large, millimeter scale probes, ultra-small microwires and silicon probes with dimensions less than 100 μm are becoming much more common, yet little is known about their insertion properties. Here, we measured insertion of a series of different size and tip shape probes with high spatial and temporal resolution for agar, *ex vivo* and *in vivo* systems. These careful mechanical insertion measurements provided a number of unexpected findings. First, the insertion force did not increase with depth *in vivo* after the pia penetration event, unlike previous *ex vivo* experiments. This implies that once through the pia layer, it would be possible to insert a probe to arbitrary depths without buckling, although it is possible that heterogeneities (e.g., white vs grey matter) could alter the trajectory of the wires.

Mechanically, both the pia penetration force and amount of tissue compression scaled linearly with probe diameter rather than cross-sectional area. The amount of tissue compression scaled as ∼ 4 times the microwire diameter, with smaller probes requiring less force and causing less tissue compression. Interestingly, no statistical difference in pia penetration force or compression was observed between flat and angle-polished wire tips, while ultra-sharp electrosharpened tips had negligible pia penetration force for all wire diameters. Once inside the tissue however, the required insertion force scaled with circumference, independent of tip shape. This indicates that the sidewalls dictate the internal insertion force, while the tip shape is more relevant for pia penetration, especially at larger probe sizes. Interestingly, rectangular 20 x 70 μm *Neuropixels* probes penetrated the pia like a 20 μm device, but inserted deeper into the tissue as a cylindrical wire with equivalent circumference.

Combined force measurements together with *in situ* epifluorescence and two-photon imaging revealed that blood vessel rupture appears to occur by a process of: (1) capture upon the probe tip, (2) elongation with further probe insertion, and (3) eventual rupture, usually not located at the probe tip. This implies that the vasculature does not fail due to encountering the tip nor a crack propagating near the tip surface, rather via attachment to the probe followed by stretching. Measurements of bleeding as a function of probe diameter discovered a size regime <25 μm where no bleeding with insertion was observed. These observations imply that by proper scaling of the probe, both tissue compression and blood vessel rupture can be avoided, which may lead to greatly improved outcomes.

The results presented here provide quantitative insights for properly scaling neural probe designs and better understanding brain mechanics during insertion of microscale devices. It is clear that the model system chosen can have a significant effect on the experimental result. From combined force and real-time measurements, we propose a tearing mechanism for blood vessel failure, which could be avoided for sufficiently small probes. Substantial additional work remains to uncover the details of these mechanisms and shed light on multiple probes inserting at one time and insertion velocity dependence. The number of surprising observations highlight the utility of basic quantitative measurements, and reveals there is still much to be discovered about interfacing artificial devices together with the brain. Design trade-offs between probe diameter, material stiffness, and buckling constraints are summarized using Euler buckling analysis in **Fig. S10**.

## Materials and Methods

### Instrumentation

We developed a high-performance mechanical measurement system. This apparatus, shown in **Fig. 1**, used *NanoMechanic*’s *iNano InForce 50* (NanoMechanics Inc, USA) indentation head as the force transducer. A displacement control protocol is used to fix the center plate of the *iNano*, while a low-noise linear actuator moves the indenter head 2.5 mm into the brain at 2 - 20 μm/s. The surgical apparatus positions the *iNano* head above the tissue, then extends the indentation head to push the microwire into the material while measuring force and displacement. A custom program was written to automate force detection of the thin water layer maintained above the hydrogel/brain surface. This was done by vibrating the nanoindenter tip and detecting a phase angle change induced by contact with the water, amplified at the interface by the capillary force pulling onto the tip. This point is referenced to be zero. The system was then programmed to insert by a user specified amount into the hydrogel/brain by a speed capped at 50 μm/s to prevent damage to the tool and assure accurate force measurements (as dictated by the internal hardware speed of the feedback loop). In a subsets of experiments (**Fig. S1a-b, S2 e-f, S7 h-i**), we used Nano Indenter XP (MTS Systems Corp., USA) with a load resolution of 0.05 μN and data acquisition frequency at 5 Hz.

### Hydrogel Preparation

An agarose 0.6 % hydrogel was made and poured into glass vials. The concentration of the hydrogel was chosen based on literature findings to best match the elastic modulus of the brain (11, 42), done previously via nanoindentation.(24) Microwires inserted into the hydrogel were not easily cleaned, and typically disposed of consequently. Attempts for cleaning were made by placing in boiling water, but subsequent insertions in fresh agarose solutions did not show repeating behavior. Using microwires freshly etched via oxygen plasma, the behavior was very consistent. During insertion experiments, water was added above the hydrogel solution to ensure hydration during the length of the test. In a subset of recording of 0.3% agarose hydrogel, *MTS Nano Indenter XP* was used (**Fig. S2 e,f**).

### Probe Fabrication

Tungsten wires of varying diameters (7.5, 15, 25, 35, 50, 80, 100 μm) were spooled, coated with Parylene C (PaC), and subsequently cut into 1” segments. Briefly, aggregates of microwires were placed into glass tubes, infiltrated with Apeizon black wax W, and subsequently polished to accomplish the desired tip angles (flat and 24º) and then released. Stainless steel rods with a 150 μm inter-diameter (ID) bore were cut via electron discharge machining (EDM) to ensure no burr existed after the cut. The parylene coating also acted to increase microwire diameter, allowing for each microwire to be coated with the needed amount to have a final OD of 140 μm, greatly reducing the deviation of each microwire as it is inserted into the 150 μm diameter bore of the stainless-steel rods. Once inserted into the rods, each wire was glued in place using EpoTek 301. Etching in oxygen plasma etched the PaC to expose a length of microwire but kept the PaC embedded in the stainless-steel tube. This method allowed for consistent minimization of angular deviation, as the nanoindentation head can only measure force in the z-axis. For electrosharpened wires, bare tungsten wires were individually submerged in 0.9 M KOH. 2V was applied between the wire and a Pt wire counter electrode using a Keithley 2600 SMU and current was recorded. Etching stopped when the submerged part of the wire broke away from the rest of the wire. This break is observed visually by use of a stereoscope and confirmed by a sharp decrease in current between the two electrodes. Sharpened tips were immediately cleaned with deionized water and isopropyl alcohol and stored for safekeeping.

Every microwire produced was imaged via SEM and documented for quality and reproducibility after manufacturing and each insertion. Cleaning of the tips post insertion was done in enzymatic soap, followed by acetone, isopropanol, ethanol, DI water, and PBS.

### Animal and ex vivo brain excision

Adult (8 to 10 month) C57BL/6J mice (JAX# 000664) were used for this study. All procedures were approved by Stanford University’s Administrative Panel on Laboratory Animal Care and Academia Sinica Institutional Animal Care and Use Committee (IACUC). Animals were anesthetized with isoflurane and decapitated. The brain was exposed and chilled with ice-cold artificial cerebrospinal fluid (aCSF) containing 125 mM NaCl, 2.5 mM KCl, 2 mM CaCl_2_, 1.25 mM NaH_2_PO_4_, 1 mM MgCl_2_, 25 mM NaHCO_3_, and 15 mM D-glucose. Freshly excised mouse brains were maintained in ice-cold aCSF, and all measurement was done within 1 hour after excision. The mechanical properties of excised brain have been found to remain constant within one hour post-mortem as long as temperature is controlled (25, 27). Repeated measurements using a 25 μm diameter wire showed deterioration of force required to insert past the pia beyond 1 hour after excision. As such, use of a quick-exchange system was developed to allow for as many insertions as possible within one hour of beginning the *ex vivo* preparation. Excised brain was stuck on a petri dish with medical grade cyanoacrylate and filled with aCSF cooled externally in an ice bath (**Fig. 1**). Surface blood vessels were avoided when possible. Prior to each insertion tests, a thin layer of aCSF was applied to the brain surface to prevent drying. Insertions were all into the motor and sensory cortex areas. Between six to eight insertions were done on each brain, and the position of each subsequent insertion was shifted by 500 μm. The nanoindenter was positioned ∼200 μm above the surface of the brain and inserted at 20 μm/s to a depth of ∼2 - 2.5 mm. The nanoindenter was pulled out rapidly (1 mm/s) to speed up each insertion and consequently how many wires we could test per brain within 1 hour after excision. Wires were cleaned with enzymatic soap and isopropyl alcohol after each insertion to ensure all tissue residue is removed. To ensure reliability of measurements, subsequent brains tests altered order of diameters used. No clear difference in pia puncture behavior was shown between the two sequential orders tested in the time allotted **Fig. S5**), suggesting the mechanical properties of the brain tissue remained stable through the recording periods (24). The large spread seen in these measurements is likely due to variations in blood vessel density or laminar structures of the mouse cortex.

### In vivo force measurements

Mice were anaesthetized with an intraperitoneal injection of 17 mg/ml ketamine and 1.7 mg/ml xylazine in saline and positioned in a stereotaxic frame. Two bone screws were placed on the skull to provide mechanical stability. A head-plate was centered to the intended surgery site on the right hemisphere and fixed to the bone screws and skull with C&B METABOND Cement (Parkell Inc., USA) or Super-Bond Cement (Sun Medical, Japan). Small (2-3 mm) craniotomies were made over the somatomotor and somatosensory cortices using standard procedures to expose the brain surface (43). Ringer’s solution was applied on the brain surface at all times to prevent tissue drying which causes bleeding. Dura mater was carefully removed with fine forceps (RS-4955, Roboz Surgical Instruments, USA) to expose the pia surface. Mice were then transferred to the experimental apparatus for force measurements under anaesthetized condition.

### In vivo two-photon and epifluorescence imaging

*In vivo* imaging were performed in adult (4-6 months old) Tg(Aldh1l1-EGFP,-DTA)D8Rth/J mice (JAX# 026033) and one Thy1-ChR2-YFP (JAX# 07612) mouse for durotomy confirmation, wherein the astrocytes express green fluorescent protein, eGFP. To reveal the blood vessels, a red fluorescent dye, Rhodamine B isothiocyanate-dextran (70kDa) solution (100 μl; 100mg/mL, Sigma, R9379), was injected through the tail vein. The mouse was mounted to an experimental apparatus on a motorized stage under the microscope (objective: XLUMPLFLN-20xW; Olympus BW51, Japan). The nanoindenter and the microwires were mounted to a micromanipulator (MP-285, Sutter, USA) with an estimated 20 degree angle (**Fig. 5a**). Two-photon imaging was performed with a custom built 2-photon laser-scanning microscope equipped with a mode-locked Ti:sapphire laser Mai Tai eHP (Spectra-Physics, USA) (44). 2-photon imaging with SHG was performed by FemtoSmart Dual (Femtonics, Hungary) equipped with a 1070 nm Fidelity-2 laser (Coherent, USA). Rhodamine B and eGFP were excited at 830 nm and 925 nm light wavelengths, respectively. Three-dimensional image stacks were taken every 50 – 100 μm advances of the microwire. Epifluorescence imaging was performed with an arc lamp and appropriate filter sets (540–580 nm for excitation; 600–640 nm for emission) to monitor the fluorescence of Rhodamine B. A CCD video camera (XC-77, Hamamatsu, Japan) was used to acquire the video at 30 frames per second.

## Acknowledgments

This study was supported by grants from the NINDS/NIH NS014861 (J.B.D. and N.A.M.), the Seed grant from Stanford Wu Tsai Neuroscience Institute (J.B.D. and N.A.M.), the GG gift fund (J.B.D.), the startup fund from Institute of Molecular Biology, Academia Sinica (Y.-W.W), the NSTC 113-2321-B-001-012 (Y.-W.W.), and the NSTC 114-2321-B-001-005 (Y.-W.W.). The authors thank Dr. Peter Cheng-Tang Pan, Mr. Ming-Cheng Lin, and Center for Nano Science and Nano Technology, National Sun Yat-sen University (NSYSU) for NanoIndention measurement (*MTS Nano Indenter XP*). The authors thank Dr. Yu-Yo Sun of the Institute of BioPharmaceutical Sciences, NSYSU, Taiwan for providing animal surgery facility. The authors thank members of the Melosh, Ding, and Wu laboratories for helpful discussions. Portions of the paper were developed from the thesis of A.O.

## Notes

### Competing Interest Statement

The authors have declared no competing interest.

### Summary of Updates

This revised version strengthens experimental validation, clarifies interpretation, expands analysis, and improves transparency and presentation throughout the manuscript. Additional control experiments were performed to validate dura free pia penetration measurements. Insertions through intact moist dura were directly compared with insertions after surgical dura removal. These measurements demonstrate substantially larger puncture forces and prolonged loading when dura is intact. Dura removal was independently confirmed using two photon second harmonic generation imaging. These controls confirm that the forces reported reflect pia penetration mechanics rather than contributions from the composite dura arachnoid pia barrier. The analysis of insertion speed effects has been expanded and clarified. Previously included Neuropixels speed dependent data have been more explicitly interpreted to distinguish changes in force displacement profile shape from penetration metrics. In addition, new slow speed measurements at 5 micrometers per second were performed using 20 and 80 micrometer cylindrical microwires to assess whether insertion rate alters diameter dependent scaling. These data show modest effects on force profiles while preserving the core mechanical scaling relationships. Additional agarose brain mimic measurements were conducted at 0.3 percent concentration. Although puncture force decreases at lower gel concentration, penetration behavior remains qualitatively distinct from brain tissue, reinforcing the conclusion that agarose does not faithfully reproduce in vivo penetration mechanics. Statistical transparency has been improved by explicitly reporting the number of insertions and animals for all main figures. Representative imaging experiments are clearly identified as such. Supplementary materials have been reorganized and expanded, including a consolidated buckling force framework incorporating crystalline silicon to clarify probe design constraints and mechanical trade offs.

