## Supplementary Information for "Ultra-sensitive measurement of brain penetration mechanics and blood vessel rupture with microscale probes"

**This PDF file includes:**

Supporting text
Figures S1 to S10
Legends for Movies S1 to S4
SI References

**Other supporting materials for this manuscript include the following:**

Movies S1 to S4

**Supporting Information Text**

**Materials considerations for designing brain-machine interfaces**

The results from this study suggest that smaller electrodes should reduce physiological damage because of reduced tissue compression. At the same time, research had found that lower material stiffness can significantly reduce traumatic response (1, 2), thus more flexible, soft probes are preferred. Yet as probes become smaller and/or more flexible they may be too compliant to bear the puncture force, $F_p$, and buckle (3). For example, flexible probes have been inserted through insertion shuttles that provide temporary mechanical stiffness to the probe. Once the probe is successfully in place, the shuttle is removed surgically, or dissolves away(4-6). Unfortunately, these shuttles must be relatively large, which may induce significant primary trauma.

The interplay between the size of the electrodes and a minimum stiffness to pierce the brain without buckling is given by the buckling force threshold, which must be larger than the force to penetrate the brain. $F_p$ was found to be primarily dependent on the diameter and tip geometry of the electrode and is independent of the material modulus. Therefore, the results from this study can predict whether a probe of a given material and size will successfully penetrate the brain before buckling. The theoretical buckling force is calculated using Euler's formula, given by:

$$\mathrm{F} = \frac{\pi^2 EI}{(KL)^2}$$

where *E* is the elastic modulus, *I* is area moment of inertia, *L* is the effective length of the wire and *K* is the effective length factor of the wire (2.0 when one end is fixed and the other end free to move laterally). **Fig. S10a** plots the buckling force as a function of probe diameter for a variety of materials commonly used as brain penetrating electrodes, assuming a cylindrical wire shape of length $l = 3\ mm$. This length was chosen because it is deep enough to reach most brain areas in mice but may need to be larger to be relevant for deeper brain regions. Plotted on top of the buckling force curves is the measured $F_p$ of flat-polished tips with wire diameter (red line). Any probe materials or sizes that lie below this line are too compliant and will buckle before penetration, while anything above can penetrate. The minimum wire diameter required for specific materials can be calculated from where the buckling force line intersects with the force to puncture (**Fig. S10b**). For example, stiff tungsten wires can be as small as 12 μm diameter, while gold has to be >20 μm without additional coatings or braces. In contrast, a compliant material like PaC needs to be >60 μm in diameter to penetrate without buckling. Conversely, these larger electrodes will induce increased initial compression in the brain, thus the benefits of softer, compliant materials must be balanced with larger diameters necessary and increased trauma.

### Figures

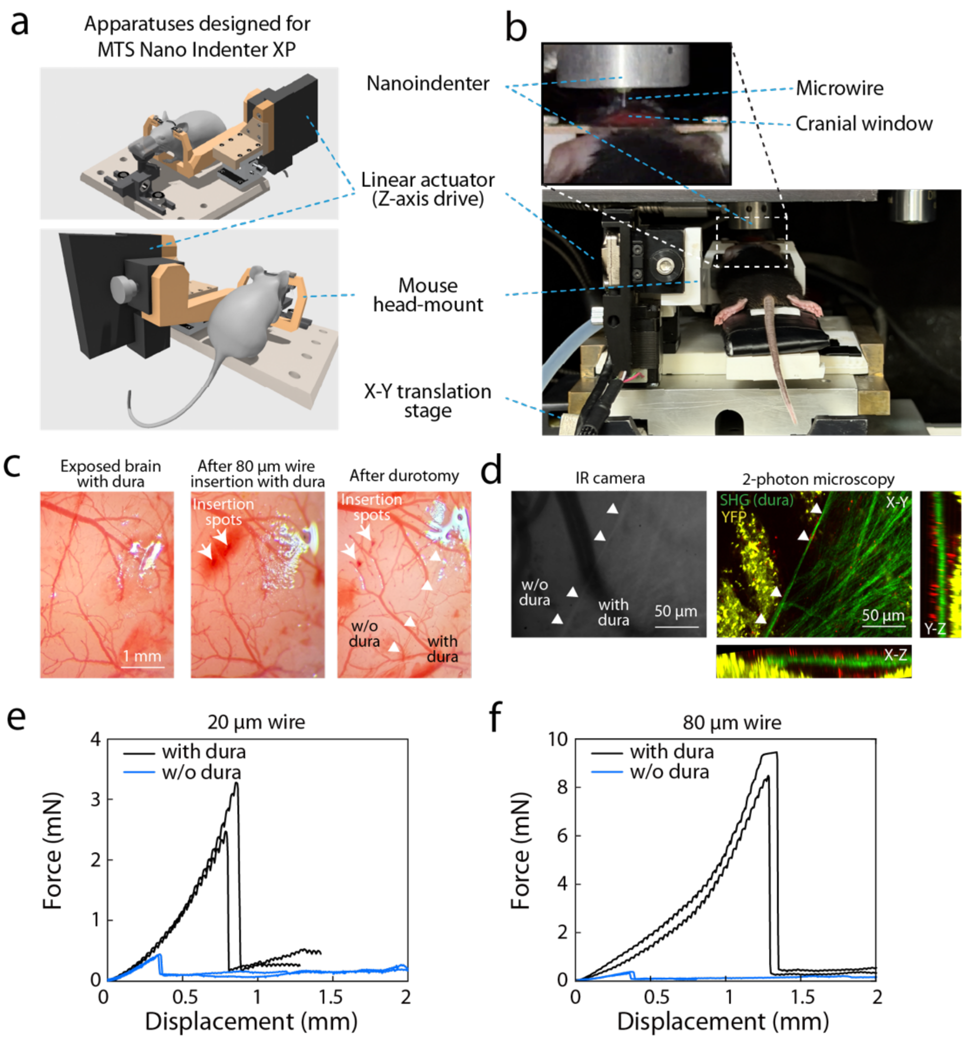


**Fig. S1. Experimental setup and control measurements for dura contribution and insertion mechanics**

**(a)** Schematic illustration of the adapted nanoindentation-based insertion system, in which an ultra-sensitive nanoindenter head (*Nano Indenter XP*) is integrated with a long-travel linear actuator to enable millimeter-scale probe insertion while recording force with nanonewton resolution. **(b)** Photograph of the *in vivo* experimental configuration showing the nanoindenter head aligned above the exposed mouse cortex mounted in a head-mounting stage under anesthesia. **(c)** Brightfield images of the cortical surface during insertion, illustrating representative conditions with intact dura mater and after surgical removal of the dura to expose the pia. Arrows mark the locations of microwire insertions and arrowheads mark the edge of the intact dura. **(d)** Left, representative light-microscopy image showing the boundary between dura-removed and dura-intact regions; arrowheads mark the edge of the intact dura. Right, two-photon microscopy image of the same region, revealing the dura via second-harmonic generation (SHG) using a 1070-nm excitation laser. **(e-f)** Representative force–displacement traces comparing insertions performed through intact dura (black) and after dura removal (light blue). Penetration through intact dura requires substantially larger forces and exhibits prolonged loading prior to rupture, whereas dura-free insertions display the characteristic pia penetration signature analyzed in the main text.

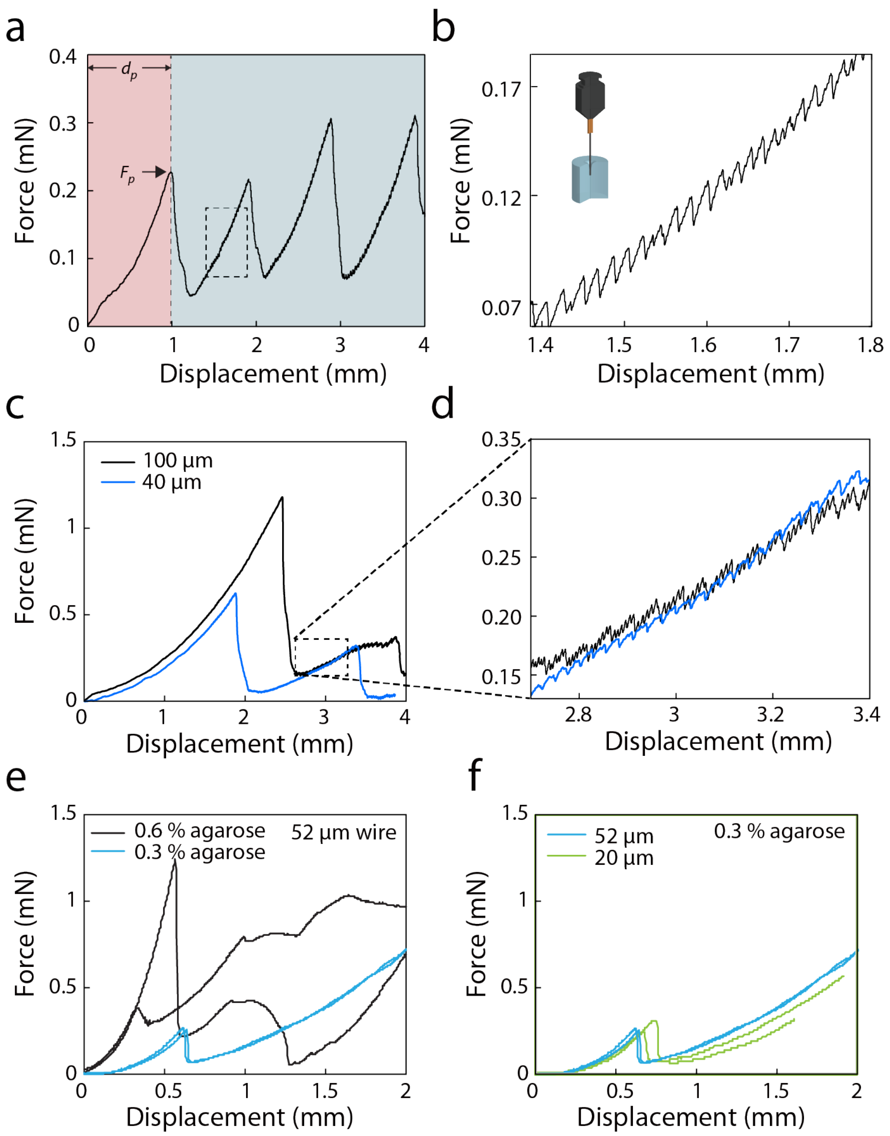


**Fig. S2. Penetration mechanics and size-dependence for agarose brain mimics**
**(a, b)** Force-displacement response for insertion of 40 µm diameter flat-polished wire with an insertion rate of 20 µm/s into 0.6% agarose brain mimic. Penetration into brain mimic and brain tissue is characterized by the sudden drop in force, wherein the puncture force ($F_p$) and displacement ($d_p$) are extracted. Large (mm scale; **a**) and small (µm scale, **(b)** sawtooth features in the force-displacement are observed post-penetration, uncharacteristic of previous measurements into brain tissue. **(c)** Representative force-displacement curves for 100 µm and 40 µm flat-polished wires into 0.6% agarose brain mimic hydrogel. **(d)** Zoom-in panel of the post-penetration phase shows differences in the saw-tooth behavior between 100 and 40 µm wires. **(e-f)** Force–displacement curves for insertion into 0.6 and 0.3% agarose, demonstrating reduced puncture force and attenuated saw-tooth behavior following penetration.

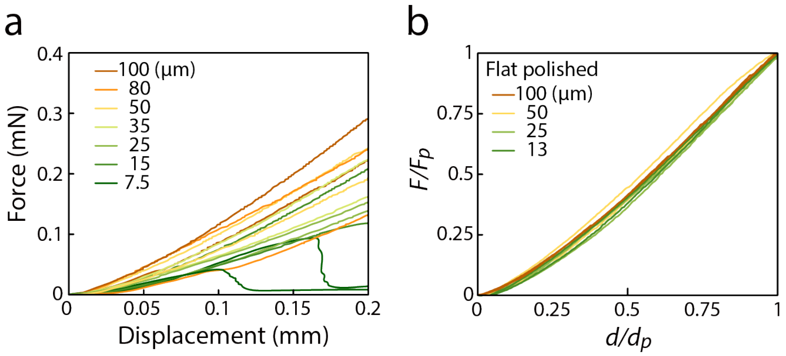


**Fig. S3. Pre-penetration curves for flat-polished wires.**

**(a)** Two representative raw traces of each size are shown. Coarse differences are found between larger and smaller wires in the force-displacement behavior pre-penetration. **(b)** Pre-penetration curves for flat-polished wires, two of each size, normalized by the force and depth at puncture. No trend with size is observed, suggesting that the underlying mechanics of indenting into brain tissue is similar between 10 – 100 µm.

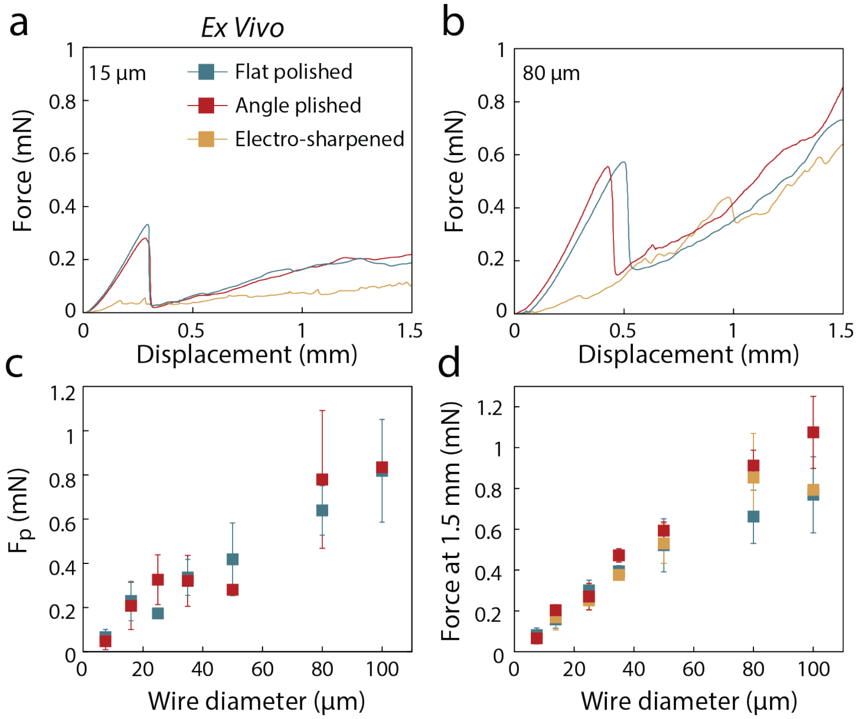


**Fig. S4. *Ex vivo* tip geometry dependence**
**(a,b)** *Ex vivo* force-displacement curves for 3 tip geometries: FP (green), AP (blue), and ES (red), for 15 µm diameter (**a**) and 80 µm diameter (**b**) microwires. (**c**) Measured puncture force ex vivo as a function of wire diameter for FP and AP wires. (**d**) *Ex vivo* force at 1.5 mm depth as a function of wire diameter for all tip geometries tested.

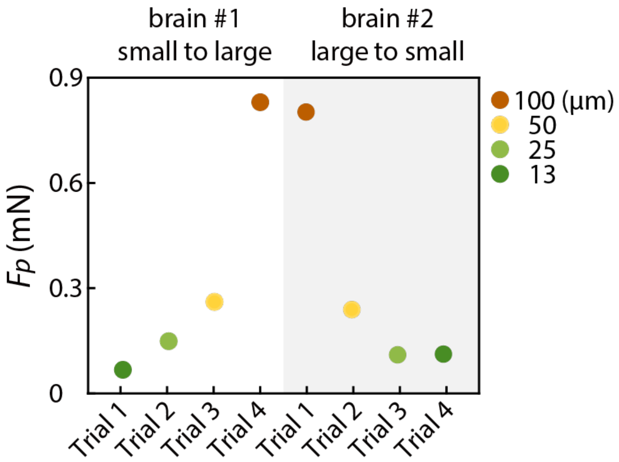


**Fig. S5. Reliability of force measurement in *ex vivo* brain tissue**
In one experiment, two brains were consecutively tested with reverse order of microwire insertion, from small to large diameters, then large to small. The puncture forces ($F_p$) are shown to be insensitive to the order in which they were tested. These results are consistent with previous work on *ex vivo* brain in different species (7).

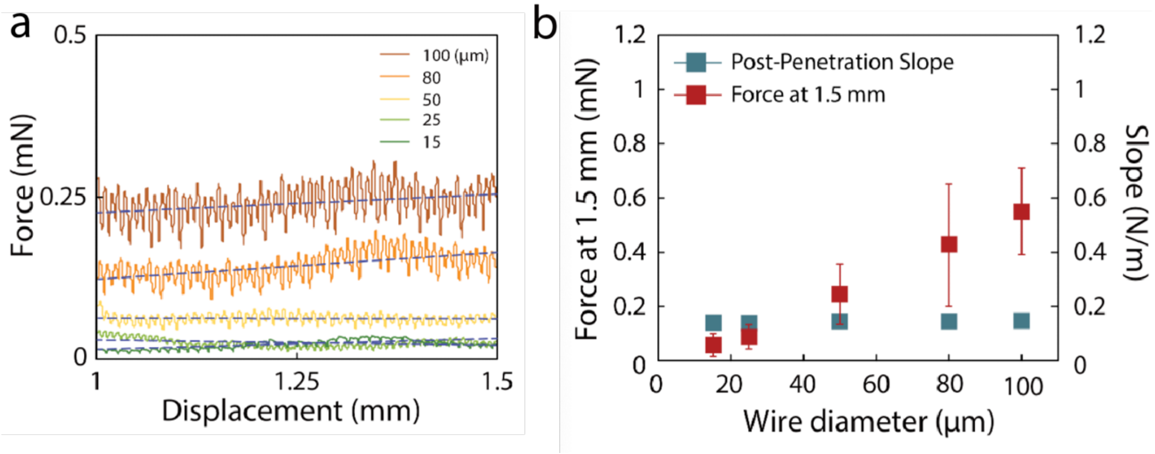


**Fig. S6. Past-penetration behavior *in vivo***
**(a)** Force-displacement behaviour of representative traces from 5 different sizes of flat-polished wires. The force oscillates due to breathing and heartbeat but doesn't increase as a function of depth. For larger microwire diameters, the oscillation amplitudes increase. **(b)** As a function of size, the constant force past-penetration scales with size (red), as do the oscillations from respiration, but the slope of this past-penetration force does not change as a function of size (blue).

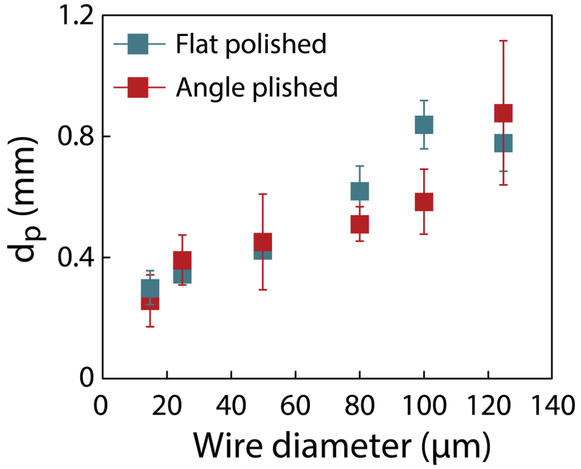


**Fig. S7.** ***In vivo*** **displacement to puncture**
Amount of brain compression before puncture as a function of wire diameter for flat and angle-polished electrodes. $d_p$ is found to scale linearly with wire diameter for both flat and angle-polished electrodes.

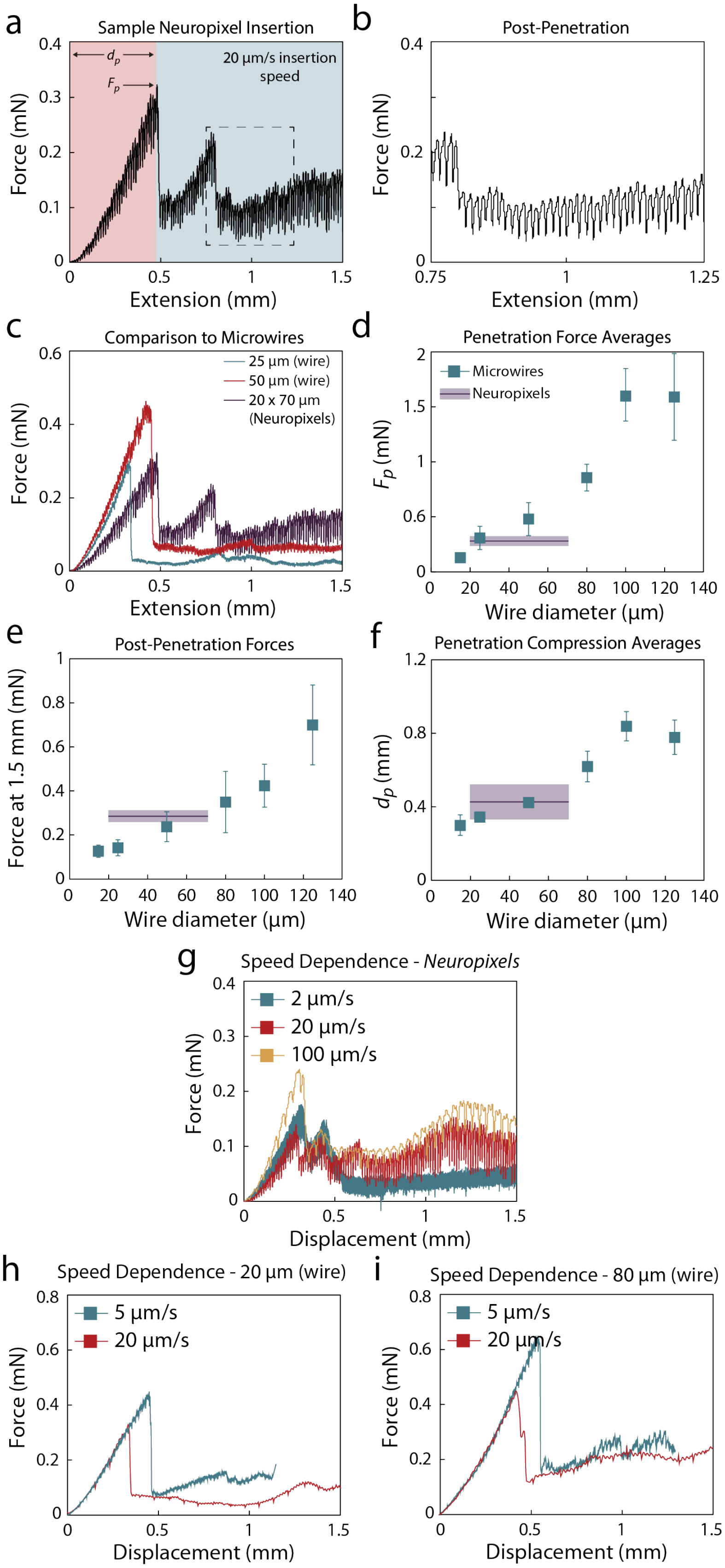
a
Sample Neuropixel Insertion
20 µm/s insertion speed
$d_p$
$F_p$
Force (mN)
Extension (mm)
b
Post-Penetration
Force (mN)
Extension (mm)
c
Comparison to Microwires
25 µm (wire)
50 µm (wire)
20 x 70 µm (Neuropixels)
Force (mN)
Extension (mm)
d
Penetration Force Averages
Microwires
Neuropixels
$F_p$ (mN)
Wire diameter (µm)
e
Post-Penetration Forces
Force at 1.5 mm (mN)
Wire diameter (µm)
f
Penetration Compression Averages
$d_p$ (mm)
Wire diameter (µm)
g
Speed Dependence - *Neuropixels*
2 µm/s
20 µm/s
100 µm/s
Force (mN)
Displacement (mm)
h
Speed Dependence - 20 µm (wire)
5 µm/s
20 µm/s
Force (mN)
Displacement (mm)
i
Speed Dependence - 80 µm (wire)
5 µm/s
20 µm/s
Force (mN)
Displacement (mm)

**Fig. S8. Neuropixels *in vivo* comparison and speed dependency**

**(a)** Sample Neuropixels inserted in vivo at 20 µm/s. Pre-penetration, **(b)** Zoom-in into the past penetration phase of the Neuropixels insertion. Oscillations due to breathing and heartbeat are consistent with measurements of cylindrical microwires. **(c)** Neuropixels insertion compared to two representative microwire insertions (50 µm in red and 25 µm in blue). The Neuropixels displays a peak penetration force similar to a 25 µm microwire **(d)**, but a compression distance and post-penetration forces closer to that of a 50 µm microwire **(e,f)**. The size values for Neuropixels in **(d-f)** are plotted between 20 – 70 µm to compare with circular microwires. **(g)** Speed dependence measurements using Neuropixels. *Fp* is slightly higher at 2 µm/s and lower at 20 µm/s; ~2x higher at 100 vs 20 µm/s), with similar penetration displacement. **(h-i)** Representative force–displacement traces for FP microwires (20 µm (**h**) and 80 µm (**i**)) inserted at 5 µm/s and 20 µm/s, illustrating modest increased *Fp* and *dp* at 5 µm/s comparing to 20 µm/s without obvious increases of post-penetration force plateaus.

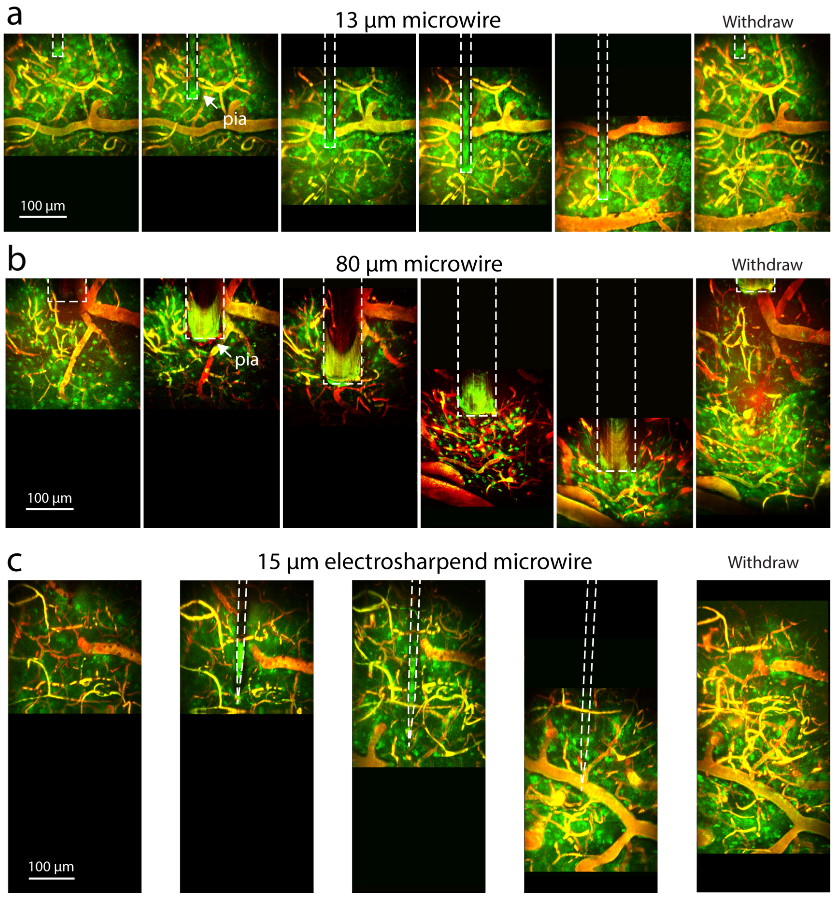


**Fig. S9. Two-photon microscopy of vascular damage**
**(a-c)** Two-photon imaging of 13 µm and 80 µm flat-polished and 15 µm electrosharpened microwire insertion, respectively. Astrocyte-GFP and BV-Rhodamine are used to stain the vasculature and astrocytes. Two-photon image stacks were taken with sequential motion of the microwire, as the imaging took several minutes to acquire, showing a few frames of the insertion process. For the 13 µm insertion, the tip of the wire drags a part of the pia surface tissue as the wire inserts. However, no disruption of the tissue is observed as the wire is driven deeper and is finally extracted. In contrast, the 80 µm wire **(b)** causes significant disruption to nearby vasculature and astrocytes. **(c)** During the insertion of a 15 µm electrosharpened microwire, no disruption is observed similar to the flat-polished small microwire **(a)**.

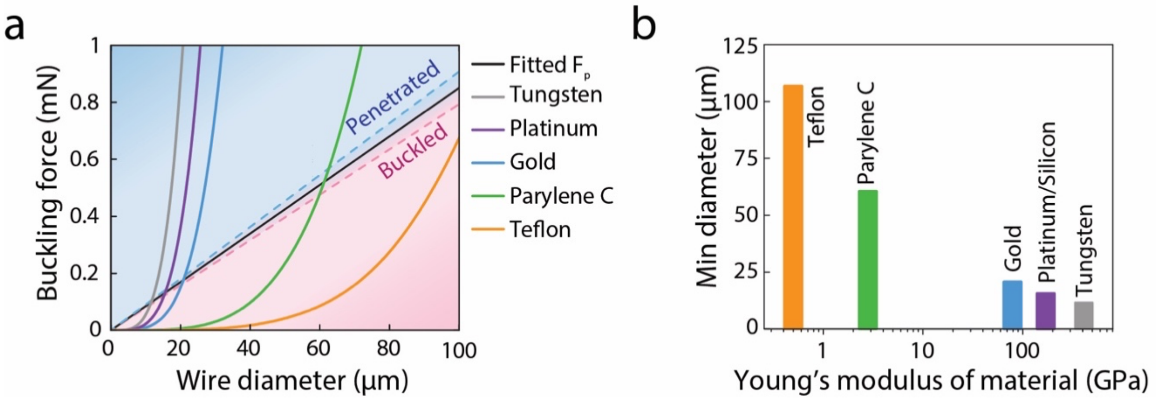


**Fig. S10. Probe material design choices**

(**a**) Buckling force curves as a function of wire diameter, calculated using Euler's buckling equation. A *K* value of 2 is used, assuming one end is rotationally and translationally fixed (distal) and the brain interfacing end is free. The length is chosen to be $L$ = 3.0 mm, a relevant length for brain-penetrating electrodes. Fitted puncture force ($F_p$) based off of flat-polished microwire dataset. Diameters in the blue region predict successful penetration into brain tissue with a wire of a specific material, while diameters in the red region are calculated to buckle before penetration. (**b**) The minimum diameter required for a specific material to penetrate into brain tissue, based off of where the theoretical buckling force intersects with the measured puncture force from (**a**).

**Movie S1. *In vivo* Insertion of 15 μm electrosharpened microwire**

Real-time video showing the insertion of a 15 μm-diameter electrosharpened tungsten microwire into the mouse motor cortex under anesthesia. The pia is gently dimpled before penetration, yet no bleeding is observed. The probe passes through the pia without visible dimpling rebound, consistent with negligible puncture force recorded in **Fig. 3 c–d** and the size-dependent "no-bleed" regime in **Fig. 4 c**. Note that the unit of puncture force in the movie is in $10^{-5}$ N.

**Movie S2. Epifluorescence imaging of 25 μm flat-polished microwire with simultaneous force measurement**

Real-time epifluorescence recording synchronized with force-displacement data (**Fig. 5 d**) during insertion of a 25 μm flat-polished tungsten microwire. The red fluorescence (Rhodamine B–dextran) labels cortical blood vessels. The movie captures the initial compression of the pia ($t_1$–$t_3$) followed by smooth penetration and progressive insertion to several hundred micrometers. Despite direct contact with small surface vessels, no dye leakage or visible hemorrhage occurs, indicating vessel displacement rather than rupture. These dynamics correspond to the "displacement zone" behavior described in the three-zone model (**Fig. 6 a–b**).

**Movie S3. Epifluorescence imaging of 80 μm flat-polished microwire with simultaneous force measurement**

Real-time epifluorescence recording of insertion of a large-diameter (80 μm) microwire, recorded concurrently with force traces shown in **Fig. 5 e**. Rhodamine B–labeled vessels collapse under pre-penetration compression ($t_1$–$t_5$), followed by multiple discrete bleeding events (#1 to #4) visible as sudden dye leakage both at the surface and deeper within the tissue. The video illustrates sequential vessel capture on the probe surface, elongation, and eventual rupture—corresponding to the "capture" mechanism of the three-zone model. Each bleeding episode aligns with saw-tooth features in the force curve.

**Movie S4. Two-photon microscopy of vascular damage**

Reconstructed 3D two-photon volumetric imaging of astrocytes (green; Aldh1l1-eGFP) and blood vessels (red; Rhodamine B–dextran) during insertion of microwires of different diameters, as shown in **Fig. S9**. *Left panel*,15 μm electrosharpened microwire: similar to the 13 μm wire, with minimal distortion and no bleeding. *Right panel*, 80 μm flat-polished microwire: significant deformation of local vasculature and astrocyte displacement, leaving a visible hole after retraction. Together these observations corroborate that tissue trauma and vascular rupture scale sharply with probe diameter and tip geometry, validating the capture–displacement–deformation model.
